# Cortical circuits implement optimal context integration

**DOI:** 10.1101/158360

**Authors:** Ramakrishnan Iyer, Stefan Mihalas

## Abstract

Neurons in the primary visual cortex (V1) predominantly respond to a patch of the visual input, their classical receptive field. These responses are modulated by the visual input in the surround [2]. This reflects the fact that features in natural scenes do not occur in isolation: lines, surfaces are generally continuous, and the surround provides context for the information in the classical receptive field. It is generally assumed that the information in the near surround is transmitted via lateral connections between neurons in the same area [2]. A series of large scale efforts have recently described the relation between lateral connectivity and visual evoked responses and found like-to-like connectivity between excitatory neurons [16, 18]. Additionally, specific cell type connectivity for inhibitory neuron types has been described [11, 31]. Current normative models of cortical function relying on sparsity [27], saliency [4] predict functional inhibition between similarly tuned neurons. What computations are consistent with the observed structure of the lateral connections between the excitatory and diverse types of inhibitory neurons?

We combined natural scene statistics [24] and mouse V1 neuron responses [7] to compute the lateral connections and computations of individual neurons which optimally integrate information from the classical receptive field with that from the surround by directly implementing Bayes rule. This increases the accuracy of representation of a natural scene under noisy conditions. We show that this network has like-to-like connectivity between excitatory neurons, similar to the observed one [16, 18, 11], and has three types of inhibition: local normalization, surround inhibition and gating of inhibition from the surround - that can be attributed to three classes of inhibitory neurons. We hypothesize that this computation: optimal integration of contextual cues with a gate to ignore context when necessary is a general property of cortical circuits, and the rules constructed for mouse V1 generalize to other areas and species.

## 1 Introduction

In recent years, several studies relating the connectivity between neurons and their in vivo activity have focused on the mouse primary visual system. It has been shown that excitatory neurons with similar orientation tuning connect to each other with higher probability than to those tuned to the orthogonal orientation [16, 18], and when they do connect, they have higher strength [6]. The connectivity rate is even higher between neurons which correlate well in response to natural scenes [16, 15]. Bock et al. [3] demonstrated, using optical physiology followed by electron microscopy (EM), a different rule for inhibitory neurons, with their inputs being nonspecific. This general principle of “like-to-like” connectivity when synaptic weights have reached their steady state is expected using a “cells which fire together, wire together” Hebbian principle [32, 33, 25].

Multiple inhibitory cell types have been described [30, 11]. While in general very complex, the relation between connectivity and cell type can still be well explained by bundling the morphological neuron types into three classes: pyramidal-targeting interneurons that target excitatory neurons and themselves, inhibitory-selective interneurons that target other classes of interneurons, and master regulators that are indiscriminate inhibitors [11]. A similar pattern of connections is observed using three inhibitory cell broad classes defined by transgenic lines [31].

How does this observed structure relate to the proposed computations of the cortical circuits?

One line of normative models (which start by postulating a computation) focuses on sparsity, the property that only a small fraction of neurons are activated by any natural image [28, 27, 29]. These models are successful at defining the classical receptive field properties of simple cells in V1. However, they predict anti-Hebbian lateral connections between excitatory neurons. These predictions remain generally true for extensions of the sparse coding models to spiking networks [40]. In networks with separate inhibitory and excitatory neurons [14], requiring sparsity leads to Hebbian connectivity between inhibitory and excitatory neurons, with no excitatory to excitatory connections.

Another very fruitful line of study has been based on Gaussian scale mixture models for the statistics of natural images [36]. These statistics have been successfully used in a model of visual processing in which the optimization is for redundancy reduction [34], having an efficient representation of the image in the absence of noise. The implementation is based on surround normalization [4], and results in predictions for V1 to act similarly to a saliency map: producing stronger representation for features which occur in isolation and are not in the vicinity of other similar features. Recent instantiations of this line of models also include a gating mechanism [5].

It has been shown in primates that contours are salient [20], and associated models rely on like-to-like connectivity [21]. Like-to-like connectivity of lateral and top-down connections have also been involved in models of perceptual organization in mid-level vision in primates [38, 26]. However these models and their extensions typically do not include knowledge of cell types involved, and there is not an exact, formal description of the computations involved.

Neurons in cortical circuits have to perform computations in the presence of noise (for a review see [8]), and possibly deal with an absence of information in the case of occlusions. Several studies on Bayesian inference focus on optimal integration of information from different sensory modalities about one feature [22].

In this study we focus on a multitude of features and their covariances, and propose that the goal of the lateral connections is to optimally integrate information from the context to minimize the influence of noise or gaps.

## 2 Results

We assume a normative model in which the lateral connections optimally integrate information from the features which are present in the surround by direct implementation of Bayes rule.

We start by assuming a simple neural code for the excitatory neurons: the firing rate of the neuron maps monotonically to the probability of the feature the neuron codes for to be in the presented image,

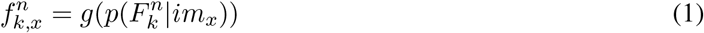

where 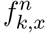 represents the firing rate of a neuron coding for feature *F_k_* at location *n* in response to image *x*, 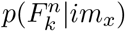 represents the probability of feature *k* being present in image *x* and g is a monotonic function. An example of such a feature is superimposed on a natural image in Figure 1a. We also assume that the sum of probabilities of all features in a patch is one, for every image, thereby implying a normalization of excitatory neuronal activities in a spatial region.

**Figure 1.**
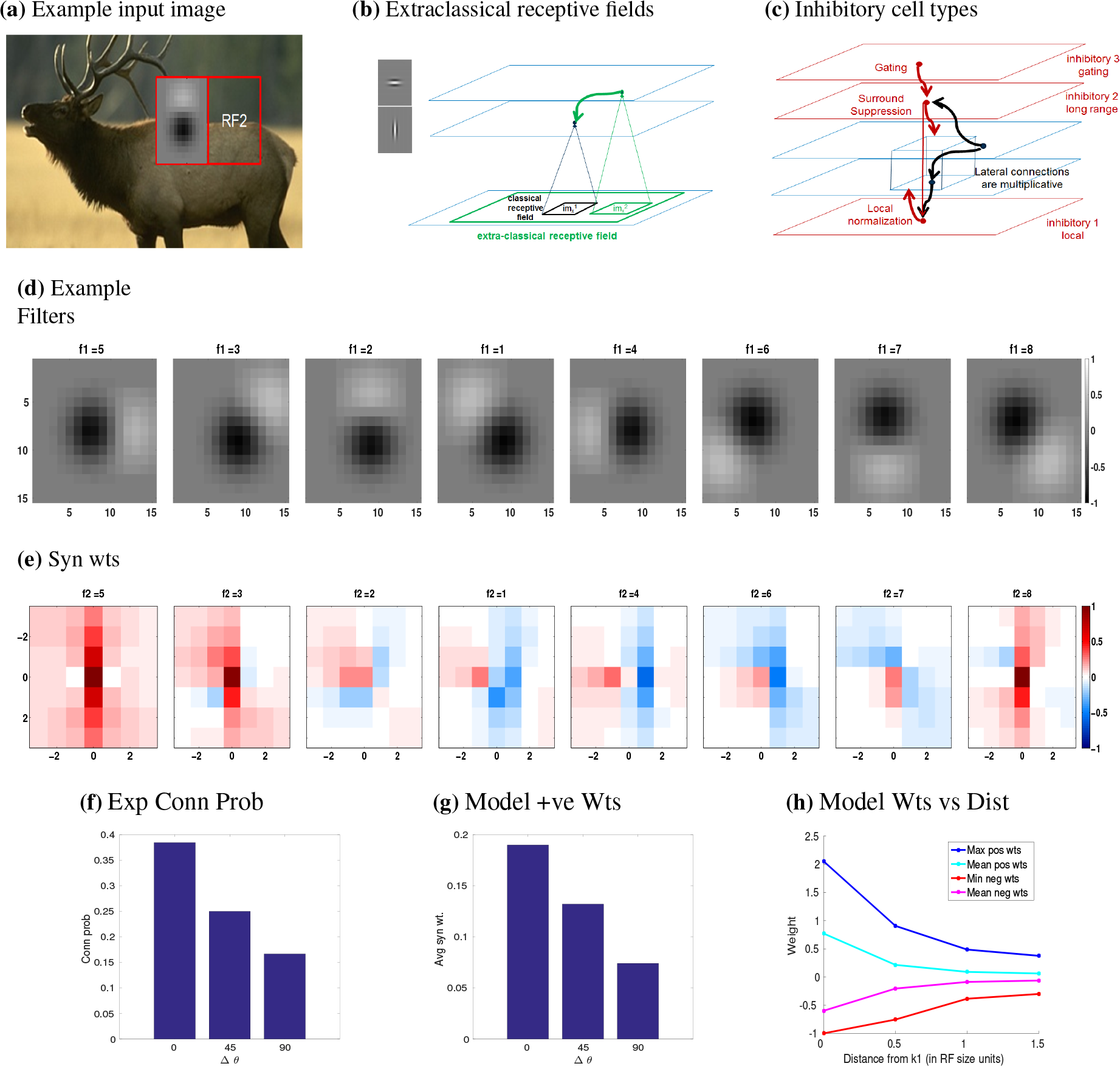
Optimal Context Integration. **a)** Sample image from the BSDS dataset, with a superimposed classical receptive field, and highlighting a surround patch (RF2) with good predictive power. **b)** Classical receptive fields are the result of feedforward connections, while extraclassical receptive fields fields are assumed to be the result of lateral or feedback connections. **c)** Three forms of inhibition are required in the model: one for local normalization, one for mediating the surround inhibition and one which gates the surround inhibition when the statistics of the surround are different from those in natural scenes. We propose that these map to classes of inhibitory neurons previously characterized [11]. **d)** A subset of basis filters were constructed using estimates of spatial receptive field (RF) sizes from in-vivo recordings. **e)** The resulting synaptic weights onto the target neuron representing the left-most filter in (d) located at position 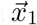 from the neurons representing filters *k*_2_ = (1,…, 8) at position 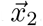. Synaptic weights were calculated at distances 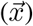 in steps of the receptive field (RF) size in each direction around *k*_1_. f) Connection probability as a function of difference in preferred orientation observed experimentally (from [16]). g) Predicted average synaptic weight as a function of difference in orientation. h) Synaptic weights fall off as distance from neurons representing filter *k*_1_ increases.

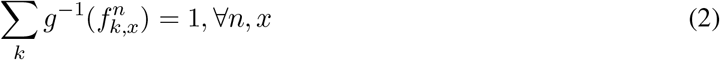

Such a normalization can be implemented by a specific inhibitory cell type, which is untuned, receives the average inputs of the excitatory neurons in its classical receptive field, and projects equally to excitatory neurons in its patch. Its connectivity matches well with the pyramidal targeting inter-neurons (PTI) category [11] and corresponds quite well to the parvalbumin-expressing inter-neurons [31]. For simplicity, we assume a linear mapping of the probability of feature presence and firing rate (*g* = 1) in the rest of the paper, as the qualitative conclusions are not dependent on this choice.

We show that a network of neurons can directly implement Bayes rule to optimally integrate the information from the surround (Methods sections 4.2 and sections 4.3). Intuitively, the activity of a neuron representing a feature is influenced by the probability that another feature is present in a surrounding patch and the statistics of co-occurrences of these features in natural scenes. The activity of a neuron representing feature *j* in patch m given image *x*, in this network can be written as:

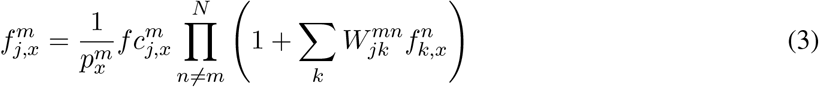

where 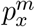 represents a normalization coefficient for patch *m* in image *x*, 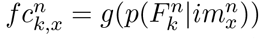 represents the classical receptive field response of the neuron and 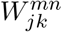 is a synaptic weight. The connection weights result from the relative probability of feature co-occurrences above the chance (Methods Equation 19) and with an identity gain function (*g* = 1), map linearly to the response rate:

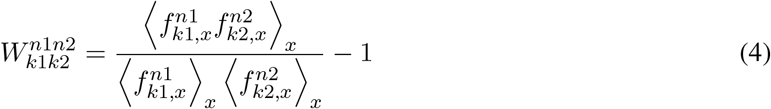

where 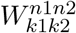 is the synaptic weight between pair (*k*_1_,*k*_2_) of filters located respectively at (*n*_1_, *n*_2_) and *x* spans the set of natural images used (in our case, the Berkeley Segmentation DataSet).

The firing rate computation requires first summing the contributions from the lateral connections corresponding to a set of neurons in a surrounding patch, and then multiplying the contributions from each set with the classical receptive field response. However, this requires quite complicated computations to be realized by individual neurons. These could potentially be realized by computations of dendritic branches. Under the assumption that the effects of the surround are small, the computation becomes simpler (Methods equation 18): all the lateral connections need to be summed, and their net influence to have a multiplicative effect on the classical receptive field input of the neuron.

To compute the predicted connectivities for mouse primary visual cortical neurons, we apply this theory to a set of simple cells. We generate a dictionary of features these neurons represent by constructing a parameterized set of models from V1 electrophysiological responses [7]. These correspond to 18 filters, one ON only, one OFF only, and 16 filters which contain both ON and OFF Gaussian subfields (Figure 1d and Methods section 4.5) at different respective relative intensities and orientations. In order to relate the activity of the neurons to the probability of a feature being present in the image as in equation 1, we convolved the image with the respective filters, rectified and then normalized the convolved output so that all values lie between 0 and 1.

We used the Berkeley Segmentation Dataset [24] to compute the probabilities of occurrences and cooccurrences of these features in natural scenes. We generated a grid of receptive fields in which the distance between gridpoints is approximately equal to the size of the classical receptive field (see Methods section 4.5). This allows the assumption that surround patches provide information which is independent of the patch in the classical receptive field (Figure 1b).

This results in a high dimensional matrix that characterizes the connection between each pair of filters at each position in the image. To reduce the dimensions and size of this matrix, we assume translational invariance: only the relative position of two filters is important. We limit it in space to three times the size of the classical receptive field (see Methods section 4.5), as the relative co-occurrence probabilities decrease significantly above this scale. The resulting connectivity matrix is 4 dimensional, with the dimensions: cell type of the source, cell type of the target and relative spatial position of the source and target in x and y directions. We present several individual 2D slices through this matrix (Figure 1e) corresponding to a specific source and target.

To compare model predictions with experimental data, we assume that one neuron in the model can be mapped to a set of neurons coding for similar features in the data. Connection weights in the model can then be interpreted as corresponding to a combination of connection probabilities and connection strength in the data. Experimentally, it has been shown that the connection probabilities and connection strength correlate well [6]. Therefore, we can compare the predicted structure to either connection strength or connection probabilities depending on which data is available.

The computation of optimal weights using equation 4 produces both positive and negative weights, with a balanced average (it should be noted that this balance is in addition to the local normalization generated by implementing Methods equation 30). Functional inhibitory connections between excitatory neurons implementing surround inhibition can be mediated by a second class of inhibitory neurons: the somatostatin positive neurons [31] and the master regulators in [11]. This is the main class of interneurons which are known to mediate surround inhibition [1].

These connections can be mapped on to a neural circuit by considering different ways of splitting the synaptic weights in equation 4 into excitatory and inhibitory terms. One possibility is to consider the excitatory term to be like-to-like and given by 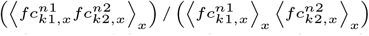. The inhibitory term, –1 is then untuned (independent of 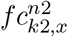) suggesting weak/non-existent orientation tuning of the interneurons mediating surround inhibition. This mapping is consistent with some experimental results [13]. For weakly correlated features, the excitatory and inhibitory terms are almost equal, leading to approximate balance. The splitting of the connection weights in to inhibitory and excitatory components in this manner results in relatively large inputs for both excitation and inhibition, resulting in a practical limitation of the spatial extent of the contextual integration window. Similar predictions with weaker overall connections would result from equating the inhibitory term with the minimum over the set of connection weights, and assigning the values above this minimum to the excitatory connections.

A different (and possibly more natural) mapping results from requiring that positive weights correspond to excitatory connections, and negative weights to inhibition. This mapping requires the inhibitory connections to be dependent on 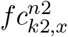 and the class of inhibitory neurons responsible for surround inhibition will be tuned to orientation. This mapping is consistent with the several experimental results [1, 23]. For the remainder of the paper, we will assume this latter mapping. This mapping predicts the excitatory connection weights as being proportional to the correlation of the responses to natural scenes, qualitatively similar to reported results [6].

The relation between excitatory neurons’ connection strength and their orientation tuning is summarized in Figure 1g. It shows a bar plot representing the mean of all positive synaptic weights for the neurons representing filters which have an orientation tuning, as a function of the relative orientation Δ*θ* between the individual subfields. The graph for excitatory connections qualitatively matches the connection probability as a function of the difference in orientation tuning (Figure 1f) reported experimentally [16]. The dependence on orientation tuning is only one of the variables and exact position and phase contribute to the computation of the synaptic weight. Note that while the theory qualitatively reproduces the dependence of synaptic weights on the difference in orientation tuning, it also predicts that sometimes cells of the same orientation but different phase (for example, panels 1 and 5 in Figure 1d) can have an inhibitory effect on each other (corresponding panels 1 and 5 in Figure 1e).

Figure 1h shows the dependence of the maximum and minimum positive and negative synaptic weights respectively onto a target neuron *k*_1_ from all neurons a fixed distance away, measured in terms of receptive field size. The plot also shows the distance dependence of the mean positive and negative synaptic weights. Assuming an exponential spatial decay, the spatial constant is *D* = 0.8 times the classical receptive field size. The spatial constant was calculated using the first two data-points in the curve for distance-dependence of the mean positive synaptic weights (cyan curve in Figure 1h). This is in accordance with previous studies [2], which suggest that the near surround extends over a range which is similar in size to the classical receptive field. Using the cortical magnification of 30*deg/mm* [10, 39], with a receptive field size of 7*deg* [7] and a spatial constant of *D* = 0.8 × receptive field size results in an average spatial constant of 187*μm*. While quite short, this distance is similar to the measured distances [19] of 114*μm* extrapolated from multipatch recordingsas measured in mouse auditory cortex.

To get insight into how optimal lateral connections might facilitate decoding of information present in neuronal activity by downstream neuronal populations, we reconstructed natural images after adding Gaussian noise (to simulate neuronal noise) to the activities of the filters (Figure 2a). We constructed maps of activities of each filter, and used the inverted filters to reconstruct the original image from all neuron activities (Methods Section 4.7 and Figure 3). We find that including such connections leads to higher reconstruction fidelity, thereby suggesting an improvement in decoding performance.

**Figure 2.**
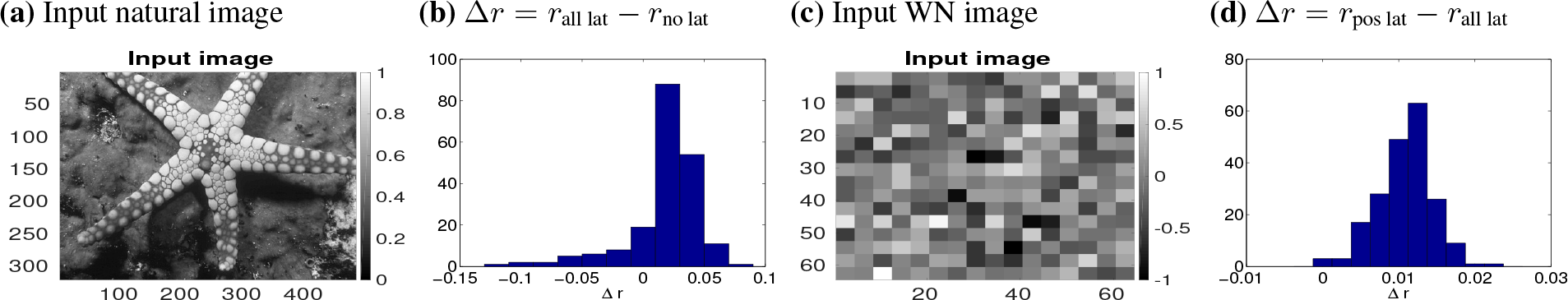
Image reconstruction from noisy representations. Inclusion of optimal lateral connections leads to higher reconstruction/decoding fidelity for natural images. For white noise images, inclusion of optimal lateral connections hurts the reconstruction, but an inhibition of the inhibitory connections (via activation of VIP neurons) leads to better reconstruction fidelity. (a) Input natural image. (b) Distribution of difference in correlation coefficients computed with all lateral connections and without lateral connections for a test set of 200 natural images from the BSDS dataset (mean *μ* = 0.0165, standard error of mean *sem* = 0.0021, ttest with p-value *p* = 2.78 × 10^-13^). (c) Unstructured white noise input image. (d) Distribution of difference in correlation coefficients computed including only lateral connections with positive weights and with all lateral connections for a set of 200 white noise images (mean *μ* = 0.0108, standard error of mean *sem* = 0.0002, ttest with p-value *p* = 1.57 × 10^-101^).

**Figure 3.**
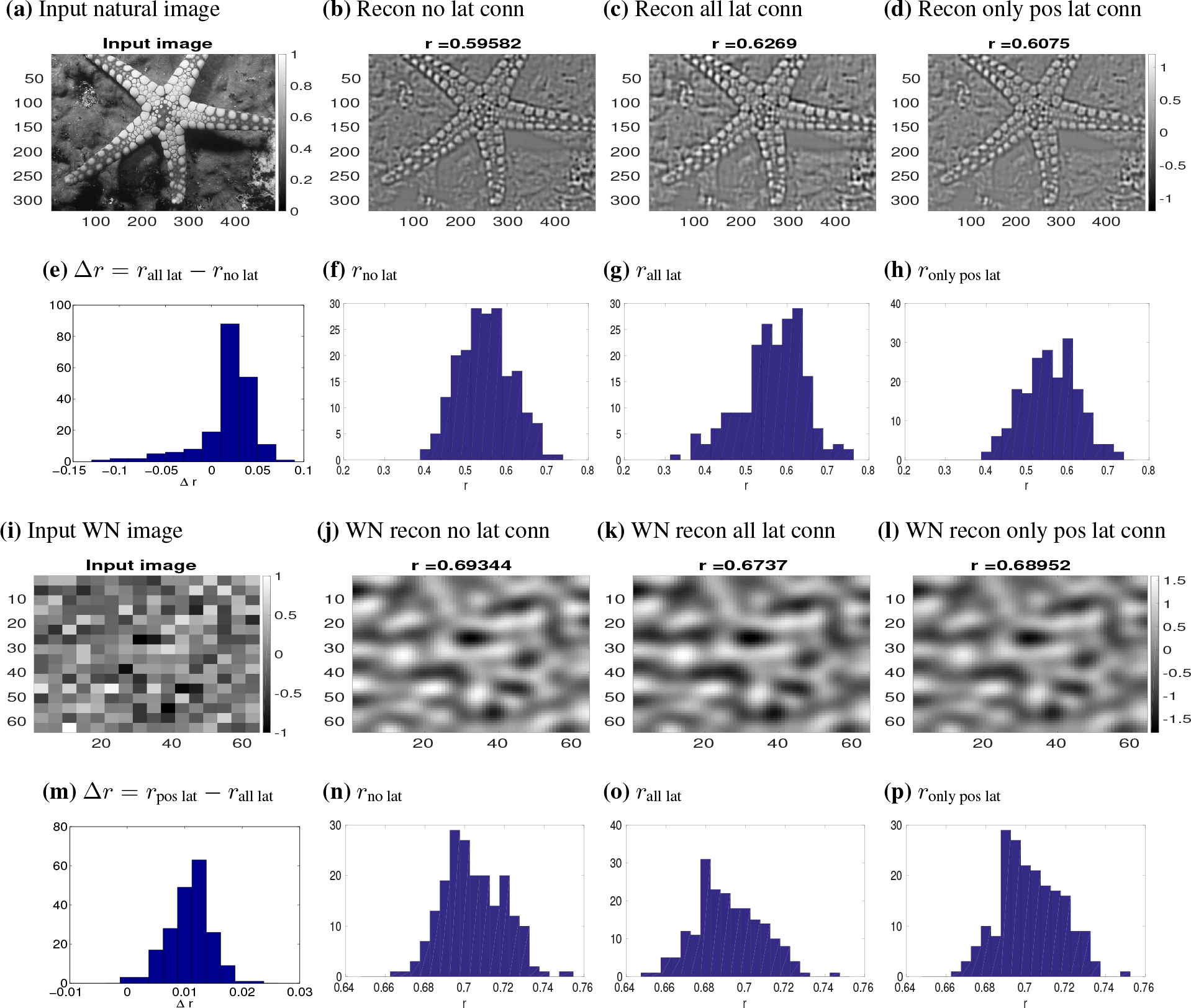
Image reconstruction from noisy representations. Inclusion of optimal lateral connections leads to higher reconstruction/decoding fidelity for natural images. **Top row:** (a) Input natural image. (b-d) Reconstructed image from inversion of activities computed (b) without lateral connections, (c) including all lateral connections, and (d) including only connections with positive weights. Correlation coefficient between respective reconstructed images and input natural image shown at the top of panels in (b-d). **Second row:** Distribution of correlation coefficients between input and reconstructed natural images from a test set (different from the training set used to compute the weights) of 200 images. (e) Distribution of difference in correlation coefficients computed with all lateral connections and without lateral connections as in columns (b) and (c) for top row (mean *μ* = 0.0165, standard error of mean *sem* = 0.0021, ttest with p-value *p* = 2.78 × 10^-13^). (f-h) Distribution of correlation coefficients for reconstructions using the same schemes as in corresponding columns of top row respectively. **Third row:** Same as in the top row, but for unstructured white noise input. Correlation coefficient between respective reconstructed images and input white-noise image shown at the top of panels in (j-1). The inclusion of optimal lateral connections hurts the reconstruction in this case, but an inhibition of the inhibitory connections (panel l) (similar to the activation of VIP neurons) allows the reconstruction to become similar to the feedforward one. **Bottom row:** Same as in second row, but for 200 white noise images. Panel (m) Distribution of difference in correlation coefficients computed including only lateral connections with positive weights and with all lateral connections as in columns (j) and (l) for third row (mean *μ* = 0.0108, standÀrd error of mean *sem* = 0.0002, ttest with p-value *p* = 1.57 × 10^-101^).

To quantify reconstruction fidelity, we calculated the Pearson correlation coefficient *r* between the input and reconstructed natural images. We find that the correlation coefficient is significantly larger for reconstruction with all lateral connections than with feedforward connections only. This can be seen in the distribution of the difference between the two correlation coefficients (Figure 2b, mean *μ* = 0.0165, standard error of mean *sem* = 0.0021, ttest with p-value *p* = 2.78 × 10^-13^). Reconstructed images and distributions of correlation coefficients for each of the respective cases can be seen in Figure 3. Although the mean of the difference between the two correlation coefficients is low, it uses lateral connections that do not require any supervised training. It is likely that repeated inclusion of information from the lateral connections through multiple layers in the visual information processing hierarchy would eventually lead to much larger differences.

When statistics of the input images do not match natural scene statistics, the optimal lateral connections can be detrimental to encoding accuracy. Could there be a mechanism to switch off the optimal context integration if the statistics do not match those of natural scenes? We test a mechanism that has an additional inhibitory gate, which inhibits the neurons implementing surround inhibition (Figure 1c). Indeed, such inhibition can be provided by neurons corresponding to the VIP [31] and inhibitory selective inter-neurons [12]. From a decoding perspective, this would imply that the reconstruction fidelity should be worse with all lateral connections included than with the positive lateral inputs alone. To test this, we used pixelated white noise images as input and reconstructed them from the activities (Methods Section 4.7 and Figure 3 therein). The white noise images consist of statistically independent patches with sizes similar to those of the subfields in our filter models. An example input image is shown in Figure 2c. In this case, we find as expected that the reconstruction based on the activity including only the positive lateral connections is better than reconstruction from activity that includes all lateral connections. This can be seen in the distribution of the difference between the two correlation coefficients (Figure 2d, mean *μ* = 0.0108, standard error of mean *sem* = 0.0002, ttest with p-value *p* = 1.57 × 10^-101^). This distribution was obtained by generating 200 instances of white noise images with different random seeds and performing the reconstructions as described above for each instance (see Methods Section 4.7). Again, reconstructed images and distributions of correlation coefficients for each of the respective cases can be seen in Figure 3

## 3 Discussion

We have presented a normative model of cortical circuit computations, in which the lateral connections among neurons serve to optimally (in a Bayesian sense) integrate information from features in the surround. Using the BSDS database of natural images [24] and classical receptive fields parameterized using mouse V1 neuron responses [7], we have computed a relation between probabilities of feature co-occurences and the synaptic weights for lateral connections which optimally integrate these features.

Mapping these connections and computations onto a network of neurons, we find that excitatory neurons show like-to-like connectivity, qualitatively in agreement with dependence of connection probability on the difference in orientation tuning that has been reported experimentally in mouse V1. Our model predicts that the strength of lateral connections between excitatory neurons should be proportional to correlations in their activity in response to natural scenes.

The mapping suggests two forms of inhibition - local normalization of excitatory neuronal activity in a patch (corresponding to classical receptive fields) and inhibition arising from the surround (extra-classical receptive fields) - that we attribute to the PV and SST-expressing interneurons respectively. When the statistics of the input image do not match those of static natural images (as can happen during when an animal is running), a gating mechanism which enables ignoring of contextual information is needed. This could be accomplished by a third form of inhibition mediated by VIP interneurons, in which there is increased activity of VIP interneurons and reduced activity of SST cells. This would effectively result in dis-inhibition of the surround, thereby allowing better processing of movement [9, 17]. When the non-linearity *g* is taken to be identity (*g* = 1), it also predicts that inputs from lateral connections should combine multiplicatively with the feedforward input to drive a neuron’s activity (thus affecting its gain).

We have also shown that including contributions from lateral connections to the feedforward activity leads to better decoding performance, when the statistics of the input image match those of natural images.

For simplicity, we have restricted ourselves to a simple basis set of filters that would extract information about oriented edges present in natural scenes, as our focus is on the biological predictions of this theory. However, the computation of the lateral connections need not be intrinsically limited to simple cells corresponding to early vision. A more extensive exploration of the computational properties of a network with such lateral connections at multiple levels is beyond the scope of the current study.

In the current study we also focus on the first order correlations. In our model, the dependence of the surround features on their own surround can be included in the second term in Methods equation 8, but it has to be partly ignored in Methods equation 13. It remains of interest to see when multiple applications of the surround are feasible. It also remains of interest if groups of multiplicative inputs, as needed for Methods equation 22 can be implemented by complex dendritic trees, or if the somewhat simpler Methods equation 23 is needed.

We have described the rules of lateral connectivity at steady state. They are similar to the connections resulting from Hebbian learning. However a full exploration of the stability of these weights under natural conditions is beyond the current work’s scope.

Models of sensory processing derived directly from statistical properties of natural images (e.g [28]) optimize for a linear basis such that the responses to natural images are as statistically independent as possible. The learned basis functions show properties resembling those of simple cell receptive fields in V1, but suggest functional inhibition between excitatory neurons. In our study, we start with a data-driven parameterization of receptive fields of mouse V1 neurons, compute synaptic weights between these neurons and demonstrate like-to-like connectivity between excitatory neurons. It is interesting to note that the formula we obtain for synaptic weights (equation 4) is formally similar to the steady state synaptic weights between excitatory and inhibitory neurons obtained in [14] from their correlation-measuring (CM) learning rule; however as it targets inhibitory neurons it serves a different function.They find that inhibitory neurons in their model decorrelate excitatory neurons and suggest a mapping of these onto PV interneurons. In contrast, we started with a proposed context integration computation for lateral connections and in mapping these to a network of neurons, suggest that PV interneurons locally normalize excitatory neuronal activity in a patch.

Taking into account sensory non-linearities, Schwartz and Simoncelli [35] developed a non-linear model of neuronal processing that removes statistical redundancy and increases independence between neuronal responses to natural stimuli via a gain control signal (computed as a weighted sum of neighboring neurons’ responses) that divisively normalizes a neuron’s response. The weights used to compute the gain control signal were chosen to maximize the independence of neuronal responses to natural stimuli. In their elegant study, Coen-Cagli et al. [4] used similar principles of redundancy reduction and natural image statistics to infer contextual interactions between the RF and surround that would lead to optimal coding of images. Using a mixture of Gaussian scale mixtures, they develop a statistical model in which the spatial heterogeneity of an image is inferred. This leads to a generalized form of divisive normalization from each RFs surround; surround suppression is engaged when the image patch falling on the RF and surround is homogeneous and is absent when it is heterogeneous. They validated this model against experimental data in a later study [5] and formulated it as a flexible normalization model that includes a gating mechanism. The framework we use here is very similar; however, we start with a different proposed role (as opposed to redundancy reduction) for lateral connections: optimal integration of context. Our framework allows us to explicitly map these interactions onto a network and suggest exact expressions for E-E excitatory synaptic weights and effective E-SST-E inhibitory synaptic weights. Our model suggests that under low contrast, occlusions or high noise conditions, it is beneficial for the center neuron to use available information from all surround neurons to provide the best possible representation it can. It is likely that both these models coexist in the brain, with the optimal integration of context being more apparent at low contrast, occlusions or other high noise conditions and redundancy reduction dominating at high contrast. It would be of particular interest to develop a model which combines the need for optimal integration of context at low contrast values to a redundancy reduction [5] at higher contrast values as a lot of the model structures are similar.

While the connectivity matrix between the predicted cell types and their functional responses is very high dimensional, we were able to test with our current quite simple model a surprisingly large number of statistics: like to like connectivity, distance dependence, three types of inhibitory neurons and their statistics of connectivity. However, continuous development of methods of functional connectivity [18, 11] will result in a finer structure and comparisons with this finer structure will probably point to the needs of including some of the terms and principles which we currently ignore: higher order correlations, more complex receptive fields, redundancy reduction, predictions of future stimuli etc.

Models which start with from a functional perspective are rare in in computational neuroscience [27, 37] and even when capturing just a part of the biological complexity, they form useful entry points for more complex models.

## 4 Methods

We first outline the computation of the optimal integration of information from nearby patches (section 4.2) under a simple neuronal code (section 4.1), in general (equation 17) and under an approximation for small contributions from each each nearby image patch (equation 18). Subsequently we show their implementation in a network (section 4.3 equations 22 and equations 23) and the relation between probabilities of feature co-occurrences and the synaptic weight which optimally integrates these features (equation 26). We subsequently describe the set of features parameterized from Durand et al. [7] (Section 4.5) and apply them to a database of natural scenes from [24].

### 4.1 Neuronal code

We assume the firing rate of neurons to represent the probability of a feature to be present in one location of the image.

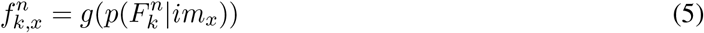

where 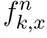 represents the firing rate of a neuron coding for feature *F*_*k*_ at location *n* in response to image *x* and g is a monotonic function. For every image and every location a normalization over features is imposed:

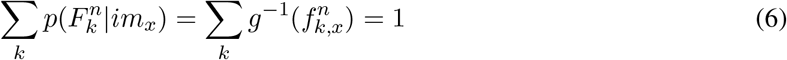

such that the sum over the probabilities of different features adds up to one. For simplicity, We will generally assume a linear mapping *g*(*x*) = *x*, however the conclusions of the study are not dependent on this choice.

### 4.2 Optimal context integration

We subdivide the image into multiple (N) sections corresponding to the sizes of classical receptive fields.

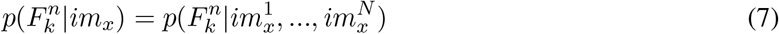

We first look at the integration of information from two patches: where patch 1 is the classical receptive field and patch 2 is the surround. The proof of optimal integration of the information from the surround will first be given for how the code for feature j, *F_j_* at location 1, 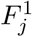, representing image x (*im_x_*), with the patch 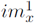 in the classical receptive field is influenced by the patch 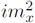 present in the extra classical receptive field.

For these two patches, equation 7 can be expended as:

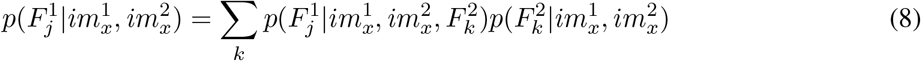

where the sum is over the *k* cells coding for the classical receptive fields in image patch 2.

We make several simplifying assumptions:

First, we assume that the only information neuron 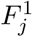 has about the patch 2 comes from the neurons with classical receptive fields in that region:

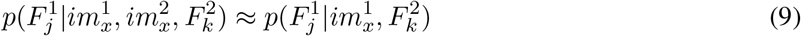

Using the approximation in equation 9, the probability of the feature 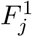 being present in *im_x_* is:

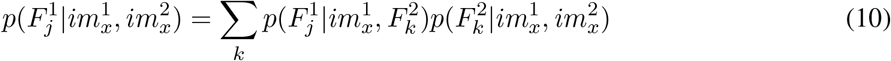

We use Bayes rule for the first term:

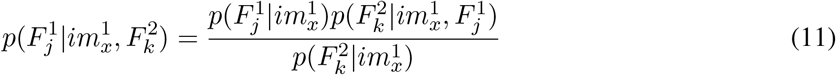

The probability described in equation 8 becomes:

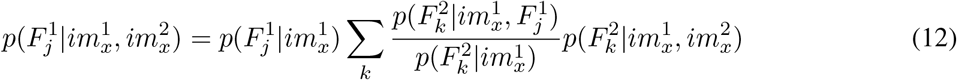

The terms 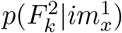 and 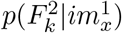 are approximately independent on the image presented. Assuming independence will allow us to implement the operation in 12 using a neuronal network. Thus for this term we make a second simplifying assumption is that:

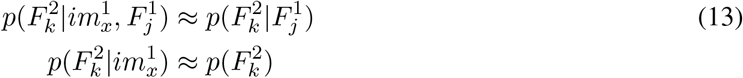

Applying this assumption on 11:

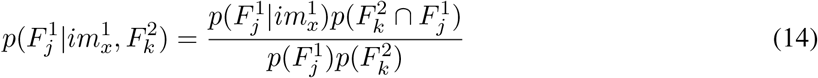

The probability described in equation 8 becomes:

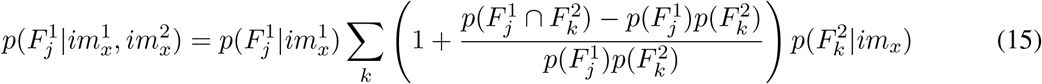

The re-arrangement of terms in 15 allows us to split the contributions as a sum of classical (feed-forward) and extra-classical (lateral) terms below, given the normalization condition 6.

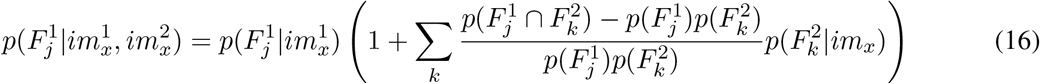

When going beyond two patches, to an arbitrary number, we use a third approximation: that each of these patches provide independent information to the neuron coding for feature *F_j_* in patch 1. The probability of feature *F_j_* in patch 1 becomes:

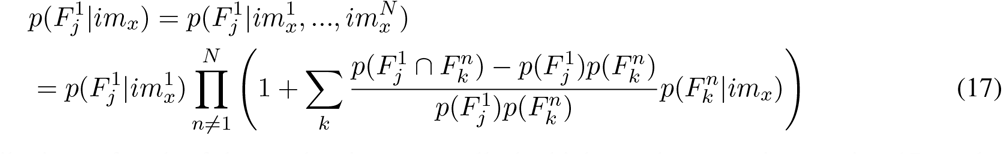

If the contributions of each of the patches is very small, the higher order terms in equation 17 can be ignored, and the probability of feature *F_j_* becomes:

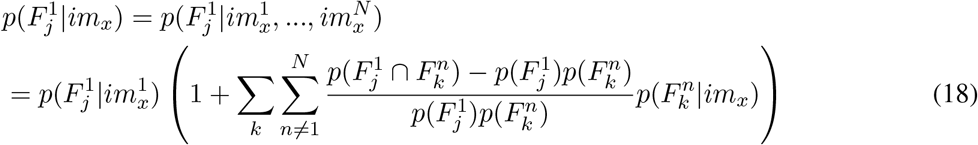

### 4.3 Cue combination under a simple neural code

Using the notation:

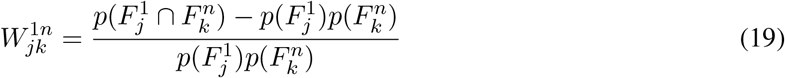

equation 17 becomes:

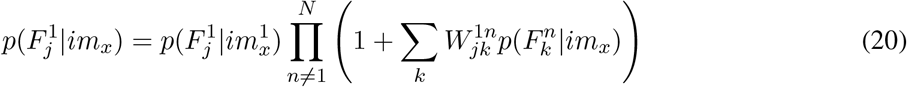

and equation 18 becomes:

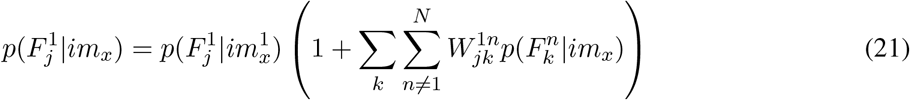

Using a simple neural code mapping *g*(*x*) = *x* in which the firing rates represent linearly the probability, the middle term can be interpreted as a synaptic weight.

The activity of the neurons representing feature j in patch 1 given image x is

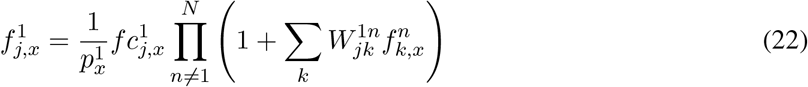

where 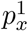 represents a normalization coefficient for patch 1 in image x and 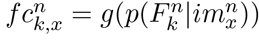 represents the classical receptive field response of the neuron. This requires the effects from the lateral connections corresponding to a group of neurons to sum, and each group to be applied multiplicatively.

Equation 18 becomes,

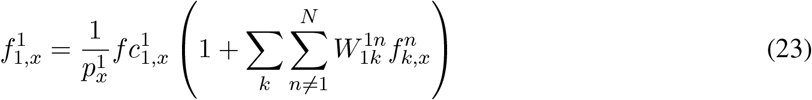

This requires the lateral connections to all be summed up and together to have a multiplicative effect.

In general, the formula for the synaptic weight,

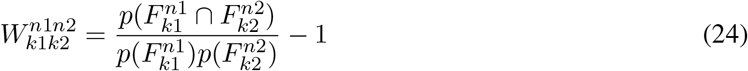

can be expressed based on average activities of the cells, when *x* spans a representative set of natural scenes, as:

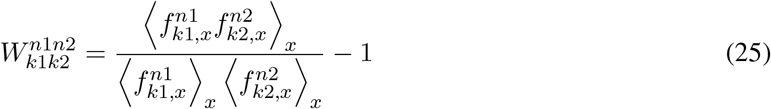

Such a set of weights can be realized using Hebbian learning in an unsupervised manner. However as an approximation, to compute these weights we used the classical receptive field responses of the cells

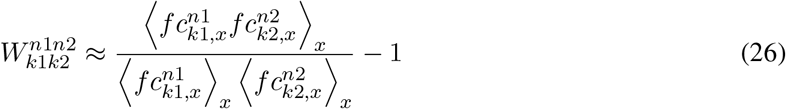

### 4.4 Normalization network

For an image patch, the we require the activity of the excitatory neurons in any patch *n* in any image *x* to be normalized:

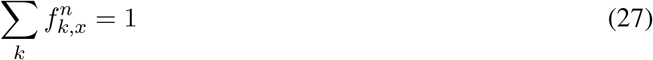

Condensing all the effect of the lateral connections into one term:

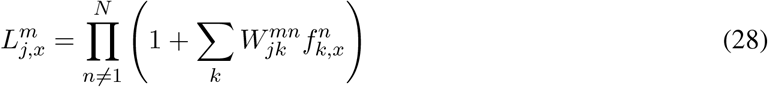

the equation for the firing rate becomes:

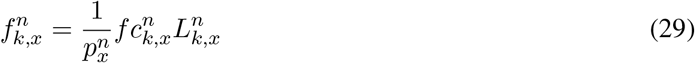

with:

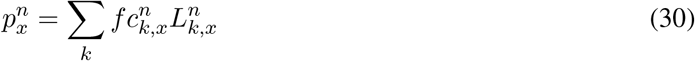

This can be implemented as a network in which a set of neurons responsible for normalization have a divisive effect on the pyramidal neurons, are patch specific (have a classical receptive field of similar size to the pyramidal neurons), are untuned, inhibit equally all the pyramidal neurons in their image patch and receive inputs equal to the average of the inputs of the pyramidal neurons in the patch.

### 4.5 Classical receptive fields parameterization

We constructed basis filters by averaging estimates of spatial receptive field (RF) sizes from 212 recorded V1 cells [7]. The basis set consisted of four classes of spatial RFs observed experimentally: ON, OFF and two versions of ON/OFF cells with the first having a stronger ON subfield and the second a stronger OFF subfield. Using the average sizes of all recorded V1 units, we modeled each subfield as a 2D Gaussian with standard deviation *σ* = 0.5 × average subfield size, which was measured to be 4.8 degrees for the OFF subfields and 4.2 degrees for the ON subfields.

The relative orientation between the two subfields for each ON/OFF class was varied from 0 to 315 degrees uniformly in steps of 45 degrees, resulting in a total of 18 filters for each spatial location. For the ON/OFF class, the distance between the centers of the two subfields was chosen to be 5 degrees (which equates to roughly 2*σ*). In accordance with data, the amplitude of the weaker subfield was chosen to be half that of the stronger subfield which was set to unity. The two subfields were then combined additively to form a receptive field whose size was 7 degrees (with size being defined as roughly equal to the distance between the two subfields plus *sigma*). A subset of these filters are shown in Figure (1c).

### 4.6 Computation of synaptic weights

We computed the synaptic weights in equation 26 as follows: Each input image *x* in the dataset was convolved with a basis filter *k* and the output was rectified and normalized so that at every spatial location *n*, the sum over all filters is unity in accordance with equation 6. An element of this rectified and normalized output corresponds to 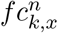.

The feature co-occurence probabilities, 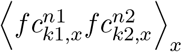 in the numerator for the synaptic weight formula in equation 26 are then computed by further pairwise convolution of the above output for each possible pair of filters in the basis set.

We computed synaptic weights 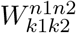 between every pair (*k*_1_, *k*_2_) of filters located respectively at (*n*_1_, *n*_2_) in the basis set described above, where x spans the set of images in the Berkeley Segmentation DataSet. We assumed translational invariance of the statistics of the image, so that

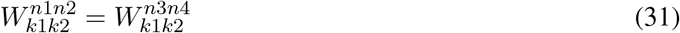

if

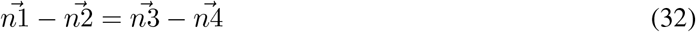

Using this assumption, for every feature/filter *k*, synaptic weights from target filters were calculated at 3 distances in steps of 0.5 × the RF size in each direction around *k*, so that we have synaptic weights on a (7 × 7) grid. Sample synaptic weights from all filters in Figure (1c) onto the first (leftmost) filter in Figure (1c) obtained using this procedure are shown in Figure (1d).

### 4.7 Reconstruction using activities and optimal synaptic weights

To get insight into how optimal lateral connections might facilitate decoding of information present in neuronal activity, we reconstructed natural images and images consisting of pixelated white noise from the respective neuronal activities. The pixelated white noise image was constructed by first generating a (16 × 16) random array and then replicating each element of this array into a (4 × 4) sub-array (each pixel is then roughly 0.5 × average subfield size), so that the entire image was effectively a (64 × 64) array. The reconstruction was performed as follows. For a given input image *x*, we calculated the effective activity 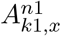 of neuron/filter *k*_1_ at location *n*_1_ using,

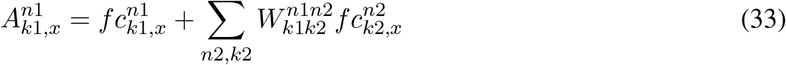

To reconstruct images from the activities, we convolved the activities computed using Equation 33 with the inverses of the filters in our basis set. Specifically, the activity 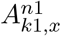 corresponding to filter *k*_1_ was convolved with the inverse of ki (obtained by flipping *k*_1_ about the horizontal and vertical axes) and the convolutions for all filters were summed together.

The reconstruction fidelity was quantified using the Pearson correlation coefficient *r* between the input and reconstructed images. The top row in Figure 3 shows the input image (panel a) and the reconstructed images based on the activity from the classical receptive field alone (panel b, *r* = 0.60), including all lateral connections (panel c, *r* = 0.63) and including only the positive lateral connections (panel d, *r* = 0.61) respectively.

The second row in Figure 2 shows the distribution of Pearson correlation coefficients between the input and the reconstructed natural images with feedforward input only (Figure 3 f), with all lateral connections (Figure 3 g) and with only the positive lateral connections (Figure 3 h) for all *test* images in the Berkeley Segmentation Dataset. The correlation coefficient is significantly larger for reconstruction with all lateral connections than with feedforward connections only. This can be seen in the distribution of the difference between the two correlation coefficients (Figure 3 e, mean *μ* = 0.0165, standard error of mean *sem* = 0.0021, ttest with p-value *p* = 2.78 × 10^-13^).

When statistics of the input images do not match natural scene statistics, the optimal lateral connections can be detrimental to encoding accuracy. To test this, we generated 200 instances of pixelated white noise images with different random seeds as input and reconstructed them from the activities as above (an example is shown in the third row in Figure 3). The white noise images consist of statistically independent patches with sizes similar to those of the subfields in our filter models. The reconstruction based on the activity from the classical receptive field alone (third row panel j, *r* = 0.69) is better than reconstruction from activity that includes all lateral connections (third row panel k, *r* = 0.67). The reconstruction from activity that includes only the positive lateral connections is also better than the one with all lateral connections included (third row panel l, *r* = 0.69) and is similar to the one with feedforward inputs alone. The beneficial effect of this gating mechanism can be seen in the distribution of the difference between the two correlation coefficients (Figure 2 d, mean *μ* = 0.0108, standard error of mean *sem* = 0.0002, ttest with p-value *p* = 1.57 × 10^-101^).

## 5 Acknowledgements

We wish to thank the founders of the Allen Institute for Brain Science, Paul G. Allen and Jody Allen, for their vision, encouragement and support, Dr. Christof Koch, Shawn Olsen, Avery Nortonsmith and Eric Shea-Brown for detailed reading of the manuscript.

## References

[1] Hillel Adesnik, William Bruns, Hiroki Taniguchi, Z Josh Huang and Massimo Scanziani. A neural circuit for spatial summation in visual cortex. Nature, 490 (7419):226-231, 10 2012

[2] Alessandra Angelucci and Bressloff Paul C.. Contribution of feedforward, lateral and feedback connections to the classical receptive field center and extra-classical receptive field surround of primate V1 neurons. In Progress in brain research, volume 154, pages 93-120. 2006

[3] Davi D Bock, Wei-Chung Allen Lee, Aaron M Kerlin, Mark L Andermann, Greg Hood, Arthur W Wetzel, Sergey Yurgenson, Edward R Soucy, Hyon Suk Kim and R Clay Reid. Network anatomy and in vivo physiology of visual cortical neurons. Nature, 471 (7337):177-82, 3 2011

[4] Ruben Coen-Cagli, Peter Dayan and Odelia Schwartz. Cortical Surround Interactions and Perceptual Salience via Natural Scene Statistics. PLoS computational biology, 8 (3):e1002405, 3 2012

[5] Ruben Coen-Cagli, Adam Kohn and Odelia Schwartz. Flexible gating of contextual influences in natural vision. Nature neuroscience, 18 (11):1648-55, 11 2015

[6] Lee Cossell, Maria Florencia Iacaruso, Dylan R Muir, Rachael Houlton, Elie N Sader, Ho Ko, Sonja B Hofer and Thomas D Mrsic-Flogel Functional organization of excitatory synaptic strength in primary visual cortex. Nature, 518 (7539):399-403, 2 2015

[7] S. Durand, R. Iyer, K. Mizuseki, S. De Vries, S. Mihalas and R.C. Reid. A comparison of visual response properties in the lateral geniculate nucleus and primary visual cortex of awake and anesthetized mice. Journal of Neuroscience, 36 (48), 2016

[8] A Aldo Faisal, Luc P J Selen and Daniel M Wolpert. Noise in the nervous system. Nature reviews. Neuroscience, 9 (4):292-303, 4 2008

[9] YuFu Jason M Tucciarone, J Sebastian Espinosa, Nengyin Sheng, Daniel P Darcy, Roger A Nicoll, Z Josh Huang and Michael P Stryker. A cortical circuit for gain control by behavioral state. Cell, 156 (6):1139-52, 3 2014

[10] M. E. Garrett, I. Nauhaus, J.H. Marshel and E.M. Callaway. Topography and Areal Organization of Mouse Visual Cortex. Journal of Neuroscience, 34 (37):12587-12600, 9 2014

[11] X. Jiang, S. Shen, C.R. Cadwell, P. Berens, F. Sinz, A.S. Ecker, S. Patel and A.S. Tolias. Principles of connectivity among morphologically defined cell types in adult neocortex. Science, 350 (6264):aac9462-aac9462, 11 2015

[12] Xiaolong Jiang, Shan Shen, Cathryn R Cadwell, Philipp Berens, Fabian Sinz, Alexander S Ecker, Saumil Patel and Andreas S Tolias. Principles of connectivity among morphologically defined cell types in adult neocortex. Science (New York, N.Y.), 350 (6264):aac9462, 11 2015

[13] Aaron M. Kerlin, Mark L. Andermann, Vladimir K. Berezovskii and R. Clay Reid. Broadly Tuned Response Properties of Diverse Inhibitory Neuron Subtypes in Mouse Visual Cortex. Neuron, 67 (5):858-871,9 2010

[14] Paul D King, Joel Zylberberg and Michael R DeWeese. Inhibitory interneurons decorrelate excitatory cells to drive sparse code formation in a spiking model of V1. The Journal of neuroscience: the official journal of the Society for Neuroscience, 33 (13):5475-5485, 3 2013

[15] Ho Ko Lee Cossell, Chiara Baragli, Jan Antolik, Claudia Clopath Sonja B Hofer and Thomas D Mrsic-Flogel. The emergence of functional microcircuits in visual cortex. Nature 496 (7443):96-100 4 2013

[16] Ho Ko Sonja B Hofer, Bruno Pichler, Katherine A Buchanan, P Jesper Sjostrom and Thomas D Mrsic-Flogel. Functional specificity of local synaptic connections in neocortical networks. Nature 473 (7345):87-91 5 2011

[17] Jung Hoon Lee and Stefan Mihalas. Visual processing mode switching regulated by VIP cells. bioRxiv, 2016

[18] Wei-Chung Allen Lee, Vincent Bonin, Michael Reed, Brett J Graham, Greg Hood, Katie Glattfelder and R Clay Reid. Anatomy and function of an excitatory network in the visual cortex. Nature, 532 (7599):370-4 4 2016

[19] Robert B. Levy and Alex D. Reyes. Spatial Profile of Excitatory and Inhibitory Synaptic Connectivity in Mouse Primary Auditory Cortex. Journal of Neuroscience, 32 (16), 2012

[20] W. Li and Charles D Gilbert. Global Contour Saliency and Local Colinear Interactions. Journal of Neurophysiology, 88 (5):2846-2856, 11 2002

[21] and Z Li.A neural model of contour integration in the primary visual cortex. Neural computation, 10 (4):903-40, 5 1998

[22] Wei Ji Ma Jeffrey M Beck, Peter E Latham and Alexandre Pouget Bayesian inference with probabilistic population codes. Nature neuroscience, 9 (11):1432-1438, 11 2006

[23] Wen-pei Ma Bao-hua Liu, Ya-tangLi Z. Josh Huang, Li I. Zhang and Huizhong W. Tao. Visual Representations by Cortical Somatostatin Inhibitory NeuronsaATSelective But with Weak and Delayed Responses. Journal of Neuroscience, 30 (43), 2010

[24] David Martin, Charless Fowlkes, Doron Tal and Jitendra Malik. A Database of Human Segmented Natural Images and its Application to Evaluating Segmentation Algorithms and Measuring Ecological Statistics

[25] Thomas Miconi, Jeffrey L McKinstry and Gerald M Edelman. Spontaneous emergence of fast attractor dynamics in a model of developing primary visual cortex. Nature communications, 7:13208, 10 2016

[26] S. Mihalas, Y. Dong, R. Von Der Heydt and E. Niebur. Mechanisms of perceptual organization provide auto-zoom and auto-localization for attention to objects. Proceedings of the National Academy of Sciences of the United States of America, 108 (18), 2011

[27] B A Olshausen and D J Field. Emergence of simple-cell receptive field properties by learning a sparse code for natural images. Nature, 381 (6583):607-609, 6 1996

[28] B A Olshausen and D J Field. Natural image statistics and efficient coding. Network (Bristol, England), 7 (2):333-339, 5 1996

[29] B A Olshausen and D J Field. Sparse coding with an overcomplete basis set: a strategy employed by V1? Vision research, 37 (23):3311-3325, 12 1997

[30] Giorgio A. Petilla Interneuron Nomenclature Group,Giorgio A Ascoli, Lidia Alonso-Nanclares, Stewart A Anderson, German Barrionuevo, Ruth Benavides-Piccione, Andreas Burkhalter, GyÁürgy Buzsáki, Bruno Cauli, Javier Defelipe, Alfonso Fairén, Dirk Feldmeyer, Gord Fishell, Yves Fregnac, Tamas F Freund, Daniel Gardner, Esther P Gardner, Jesse H Goldberg, Moritz Helmstaedter, Shaul Hestrin, Fuyuki Karube, ZoltÁan F Kisvárday, Bertrand Lambolez, David A Lewis, Oscar Marin, Henry Markram, Alberto Muñoz, Adam Packer, Carl C H Petersen, Kathleen S Rockland, Jean Rossier, Bernardo Rudy, Peter Somogyi, Jochen F Staiger, Gabor Tamas, Alex M Thomson, Maria Toledo-Rodriguez, Yun Wang, David C West and Rafael Yuste. Petilla terminology: nomenclature of features of GABAergic interneurons of the cerebral cortex. Nature reviews. Neuroscience, 9 (7):557-568,7 2008

[31] Carsten K Pfeffer, Mingshan Xue, Miao He, Z Josh Huang and Massimo Scanziani. Inhibition of inhibition in visual cortex: the logic of connections between molecularly distinct interneurons. Nature neuroscience, 16 (8):1068-1076, 8 2013

[32] Sadra Sadeh,Claudia Clopath and Stefan Rotter. Emergence of Functional Specificity in Balanced Networks with Synaptic Plasticity. PLoS computational biology, 11 (6):e1004307, 6 2015

[33] Sadra Sadeh,Claudia Clopath and Stefan Rotter. Processing of Feature Selectivity in Cortical Networks with Specific Connectivity. PloS one, 10 (6):e0127547, 6 2015

[34] Odelia Schwartz, Terrence J. Sejnowski and Peter Dayan. Soft Mixer Assignment in a Hierarchical Generative Model of Natural Scene Statistics. Neural Computation, 18 (11):2680-2718, 11 2006

[35] Odelia Schwartz and Eero P. Simoncelli. Natural signal statistics and sensory gain control. Nature Neuroscience, 4 (8):819-825, 8 2001

[36] Martin J. Wainwright and Eero P. Simoncelli. Scale Mixtures of Gaussians and the Statistics of Natural Images, 2000

[37] and Daniel Yamins. Performance-optimized hierarchical models predict neural responses in higher visual cortex. Proceedings of the National Academy of Sciences of the United States of America, 2014

[38] and Li Zhaoping. Border Ownership from Intracortical Interactions in Visual Area V2. Neuron, 47 (1):143-153,7 2005

[39] Jun Zhuang, Lydia Ng, Derric Williams, Matthew Valley, Yang Li, Marina Garrett and Jack Waters. An extended retinotopic map of mouse cortex. eLife, 6, 1 2017

[40] Joel Zylberberg, Jason Timothy Murphy and Michael Robert DeWeese. A sparse coding model with synaptically local plasticity and spiking neurons can account for the diverse shapes of V1 simple cell receptive fields. PLoS computational biology, 7 (10):e1002250, 10 2011.

